# Kernel Dimensionality Has Limited Impact on Generalizability of CNN-Based Neural Drive Estimation from HD-sEMG

**DOI:** 10.64898/2026.03.20.712696

**Authors:** Jirui Fu, Helen J. Huang, Yue Wen

**Author notes:** (Corresponding author: Yue Wen).

## Abstract

Convolutional neural networks (CNNs) have been widely used to estimate neural drive from high-density surface electromyography (HD-sEMG) signals in neural machine interfaces owing to their real-time capability. Depending on kernel dimensionality (1D, 2D, or 3D), CNNs can extract temporal, spatial, or spatiotemporal features. Given that motor unit action potentials propagate across both space and time, architectures that exploit spatial features may offer advantages for neural drive estimation. Despite the potential importance of kernel dimensionality, its influence on neural drive estimation remains poorly understood. Existing studies have mainly evaluated CNN generalizability across participants, contraction intensities, or muscles within the same HD-sEMG dataset, while computational efficiency has seldom been considered. As a result, it remains unclear whether different kernel dimensionalities affect cross-dataset generalizability and computational efficiency. In this study, we implemented three CNN architectures—differing only in kernel dimensionality— to investigate whether exploiting the spatial and spatiotemporal features of motor unit action potentials improves the generalizability and computational efficiency of neural drive estimation from HD-sEMG recorded during lower-limb isometric contractions. We trained the CNNs on one HD-sEMG dataset and evaluated them, without retraining, on two independent, unseen datasets recorded from different participants, sessions, and protocols—one spanning three contraction intensities and the other three muscles. All three architectures are generalized to both unseen datasets. The 2D and 3D CNNs marginally outperformed the 1D CNN with a 0.2% increase in R, while the 3D CNN showed no advantage over the 2D CNN. Computational efficiency depended on kernel dimensionality in a platform-dependent manner. On the CPU, the 3D CNN showed the slowest inference time, which was 2× slower than the 2D and 1D CNN, owing to the higher arithmetic cost of its spatiotemporal convolutions. On the GPU, all three architectures achieved similar inference times of about 1.36 ms/sample. These findings indicate that increased architectural complexity of CNN does not improve generalizability for neural drive estimation, and that a 2D CNN offers the best balance of accuracy and efficiency for a reliable, deployable CNN-based neural drive estimator—particularly on CPU-only or resource-constrained platforms.

## I Introduction

The neural drive to muscles estimated by surface electromyography (sEMG) has been widely used as a myoelectric controller in the applications of neural machine interface (NMI), such as estimating the muscle force in stroke survivors to control the rehabilitative device [1] and controlling the movement of a robotic hand [2]. Unlike the myoelectric control system driven by raw sEMG signals, which require frequent recalibration across participants, muscles, and contraction intensities [3], [4], the neural drive is more robust against these variations, it originates from the transformation of synaptic input from the central nervous system to recruited motor units which generates motor unit action potentials (MUAPs), and therefore reflects the neural command to drive the muscle [5], [6]. However, the MUAPs that propagate from recruited motor units are superimposed and distorted by the skin and subcutaneous tissue, which significantly impairs the accuracy of neural drive estimates from sEMG. To improve accuracy, high-density sEMG (HD-sEMG)—a variant of sEMG recorded with an array of closely spaced electrodes—has been introduced. The HD-sEMG electrodes can capture both the temporal and spatial features of MUAPs, which provide richer information for estimating neural drive [7].

The standard methods for estimating neural drive from HDsEMG rely on blind source separation (BSS), which uses offline BSS algorithms—such as FastICA-based methods [8], [9] or convolution kernel compensation [10]—to optimize a separation matrix that decomposes HD-sEMG into individual motor unit spike trains for neural drive estimation [5], [6]. However, because the separation matrix is fixed at the time of decomposition, BSS-based methods cannot readily be applied in real-time NMI applications: MUAP features change over time due to factors such as muscle fatigue and rotation of motor units [11], [12], which invalidate the fixed separation matrix and degrade estimation. Although efforts have been made to extend BSS to real-time use—for example, by periodically updating the separation matrix to track MUAP changes during a task [13], [14]—these approaches incur additional computational cost and introduce extra latency. To address these limitations, deep convolutional neural networks (CNNs), originally developed for image and video recognition, have been adopted as an alternative approach for neural drive estimation. Using collected HD-sEMG data for training, the CNNs optimize convolutional kernels with different dimensionalities to extract temporal (1D), spatial (2D), or spatiotemporal (3D) MUAP features. Several recent studies have demonstrated the capability of CNNs to estimate neural drive in real-time by using the pre-collected HD-sEMG signal and ground truth neural drive estimated by BSS algorithm to train CNNs. For example, [15] applied a 2D CNN to estimate neural drive and predict finger force, [16] introduced a multi-head 1D CNN to estimate neural drive, and [17] studied the impact of sliding window and step size on a 3D CNN for neural drive estimation.

However, HD-sEMG signals vary substantially across the conditions under which they are recorded—including contraction intensity, muscle, participant, and recording session— owing to anatomical, neural, and methodological factors [18], [19]. Therefore, in NMI applications, a CNN trained on one HD-sEMG dataset may fail to generalize to another dataset recorded under different conditions. To mitigate this challenge, some studies implemented transfer learning, using a small portion of the HD-sEMG signals collected under different conditions to fine-tune CNN models, thereby improving their generalizability across these conditions. For example, [20] and [21] used transfer learning to improve the generalizability of CNNs across different sessions and contraction intensities. However, this approach still requires collecting condition-specific HD-sEMG data and additional model fine-tuning, limiting the immediate applicability of CNNs and increasing computational and time requirements prior to deployment. Alternatively, other studies have focused on improving the inherent ability of CNNs to generalize across variations in HD-sEMG without retraining the model. For example, [15] reported generalization across movements but not across participants, and [22], [23] demonstrated generalization across intensities, muscles, and participants. However, the reported generalizability was evaluated under partially familiar recording conditions, in which at least one factor varied between training and evaluation while other factors had already been observed during training. Consequently, the models were not assessed on data in which all relevant sources of variation were simultaneously unseen.

In practical NMI applications, however, deployment data are almost always different from the training data and may originate from participants, muscles, contraction intensities, recording sessions, or movements unseen during training. A CNN suitable for robust NMI must therefore generalize to an independent HD-sEMG dataset collected under entirely unseen conditions, yet this capability has not been systematically evaluated. Moreover, the factors governing CNN generalizability remain largely unexplored. Because the spatial features of MUAPs captured by HD-sEMG have been shown to generalize across individuals [24], and because these spatial features can be extracted by 2D (spatial) and 3D (spatiotemporal) convolutional kernels, kernel dimensionality may influence CNN generalizability. However, this possibility has not been investigated, and the role of kernel dimensionality in determining CNN generalizability remains unknown.

Therefore, the objective of this study is to examine how CNN kernel dimensionality—corresponding to temporal (1D), spatial (2D), or spatiotemporal (3D) feature extraction— influences the generalizability and efficiency of decoding the neural drive, quantified by the cumulative spike train (CST), from the unseen HD-sEMG dataset in real time. We hypothesized that higher-dimensional kernels would enhance generalizability by exploiting the spatial or spatiotemporal features of HD-sEMG signals but at the cost of greater computational complexity; accordingly, we expected the 3D CNN to achieve the highest generalizability and the 1D CNN to offer the greatest computational efficiency. We focused on estimating neural drive from HD-EMG recorded from lower-limb muscles and used the BSS algorithm to decompose HDsEMG into motor unit spike trains, which were accumulated into the CST, serving as the “ground-truth” neural drive for training and evaluating the CNN models. We then adopted a unified CNN architecture from our previous work [17], [23] and trained three models with identical network structures but different kernel dimensionalities. We evaluated their generalizability and computational efficiency on two independent HDsEMG datasets, each collected under a different experimental protocol and from different muscles, contraction intensities, and subjects, with one dataset spanning multiple contraction intensities within a single muscle and the other spanning multiple muscles at a single contraction intensity.

## II Methods

### A. HD-sEMG Datasets

We used three HD-sEMG datasets in this study (Fig. 1), each collected under a distinct experimental protocol: one for training and validating deep CNNs (training dataset), and two for evaluating the generalizability of the trained models (evaluation datasets). The training dataset was collected from multiple participants and muscles under a single contraction intensity across different days. The two evaluation datasets were collected on different days from non-overlapping participant groups: one from a single muscle across three contraction intensities (cross-intensity dataset), and the other from three muscles at a single contraction intensity (cross-muscle dataset). Consequently, both evaluation datasets enabled the evaluation of cross-intensity, cross-muscle, and cross-participant generalizability of deep CNNs. Across all three datasets, HD-sEMG signals were recorded during a commonly used task in this field—repeated isometric lower-limb contractions following a trapezoidal intensity trajectory, performed on a dynamometer (Biodex, USA). Signals were sampled at 2048 Hz using a two-dimensional adhesive HD-sEMG grid of 13 × 5 gold-coated electrodes (8 mm inter-electrode distance, with one missing corner electrode; ELSCH064NM2, OT Bioelettronica, Italy). The recording protocol for each dataset was approved by the relevant institutional research ethics committee, and all procedures complied with the Declaration of Helsinki.

**Fig. 1:**
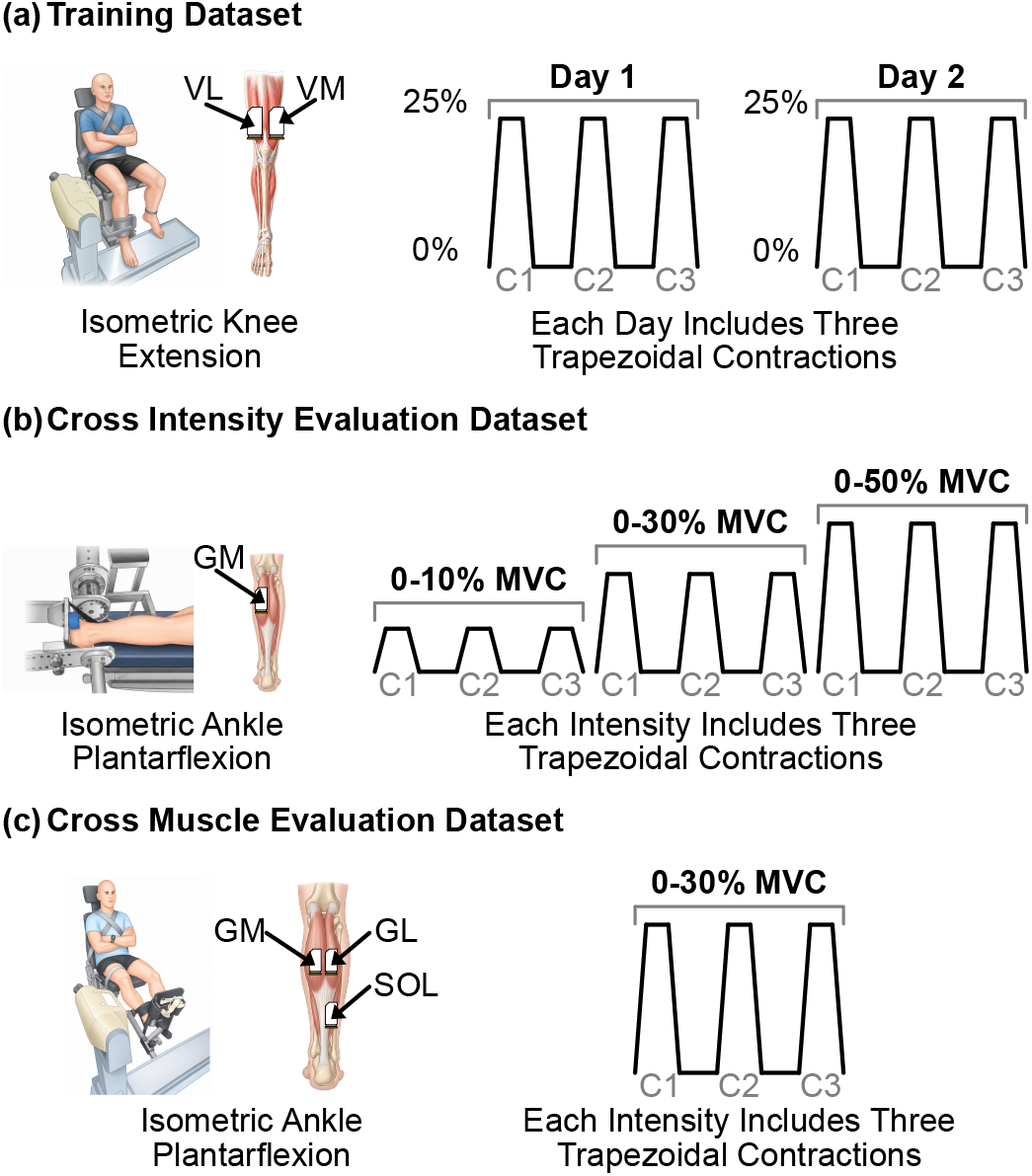
Experimental protocols of the datasets used to train and evaluate the deep CNN models. (a) The training dataset was recorded from the VL and VM muscles during isometric knee extension in five participants on two separate days at 25% MVC. (b) The cross-intensity evaluation dataset was recorded during isometric ankle plantarflexion across three intensities (10%, 30%, and 50% MVC) from GM muscle in seven different participants on a single day. (c) The cross-muscle evaluation dataset was also recorded during isometric ankle plantarflexion across three muscles (GM, GL, and SOL) in six different participants on a single day at 30% MVC.

#### 1) Training Dataset

Five young healthy participants were enrolled in this dataset (29.4±7.9 years; height 180±7 cm; body mass 76±8 kg) with no history of lower-limb injury in the past 6 months [25]. The protocol was approved by the institutional research ethics committee ‘Comité de protection des personnes Île-de-France XI’ (CPP-MIP-013). As shown in Fig. 1(a), HD-sEMG signals were recorded in two sessions separated by approximately 20 months. During each session, signals were collected from the vastus lateralis (VL) and vastus medialis (VM) muscles at 25% MVC. Participants were instructed to perform three repeated isometric knee extensions (0^*°*^–80^*°*^, where 0^*°*^ denotes full extension), with each extension following a trapezoidal intensity trajectory (ramp-up, plateau, ramp-down), and the plateau held at target contraction intensity.

#### 2) Cross-Intensity Evaluation Dataset

Seven healthy participants, independent of the training dataset (30±6 years; height 183±6 cm; body mass 74±8 kg) with no history of lower-limb injury in the past 6 months, were included in this dataset [26]. The protocol was approved by the institutional research ethics committee of the University of Queensland (No. 2013001448). As shown in Fig. 1(b), the dataset comprised HD-sEMG signals recorded from the gastrocnemius medialis (GM) muscle on a single day. Participants performed three repeated isometric ankle plantarflexions (0^*°*^–10^*°*^) at three intensities (10%, 30%, and 50% MVC), following the same trapezoidal intensity trajectory as in the training dataset, but with different ramp-up, plateau, and ramp-down durations.

#### 3) Cross-Muscle Evaluation Dataset

The cross-muscle evaluation dataset comprised six healthy participants independent of the training and cross-intensity evaluation datasets (28.8±7.0 years; height 181.3±7.1 cm; body mass 74.1±7.5 kg) with no history of lower-limb injury in the past 6 months [27]. The protocol was approved by the institutional research ethics committee ‘Comité de protection des personnes Île-deFrance XI’ (CPP-MIP-012). As shown in Fig. 1(c), the HDsEMG signals were recorded from the gastrocnemius medialis (GM), gastrocnemius lateralis (GL), and soleus (SOL) muscles in a single-day session. Participants performed isometric ankle plantarflexion following a trapezoidal intensity trajectory, with the plateau maintained at 30% MVC.

### B. Architecture of Deep CNN Models

We developed a deep CNN model to estimate the neural drive from the HD-sEMG signal. As shown in Fig. 2, the model consists of two serially connected convolutional blocks, followed by a flatten layer and a multi-head output module that decodes the neural drive as a cumulative spike train (CST). Each convolutional block contains two convolutional layers— each followed by a batch normalization layer and a ReLU activation function—and concludes with a max-pooling and a dropout layer. The multi-head module comprises four parallel heads; each head passes the flattened features through a fully connected layer with ReLU activation function and dropout layer, followed by a second fully connected layer with a sigmoid activation function, producing a binary output. The four heads encode the CST ordinally: head *k* predicts whether the CST is at least *k* (i.e., CST ≥ *k*).

**Fig. 2:**
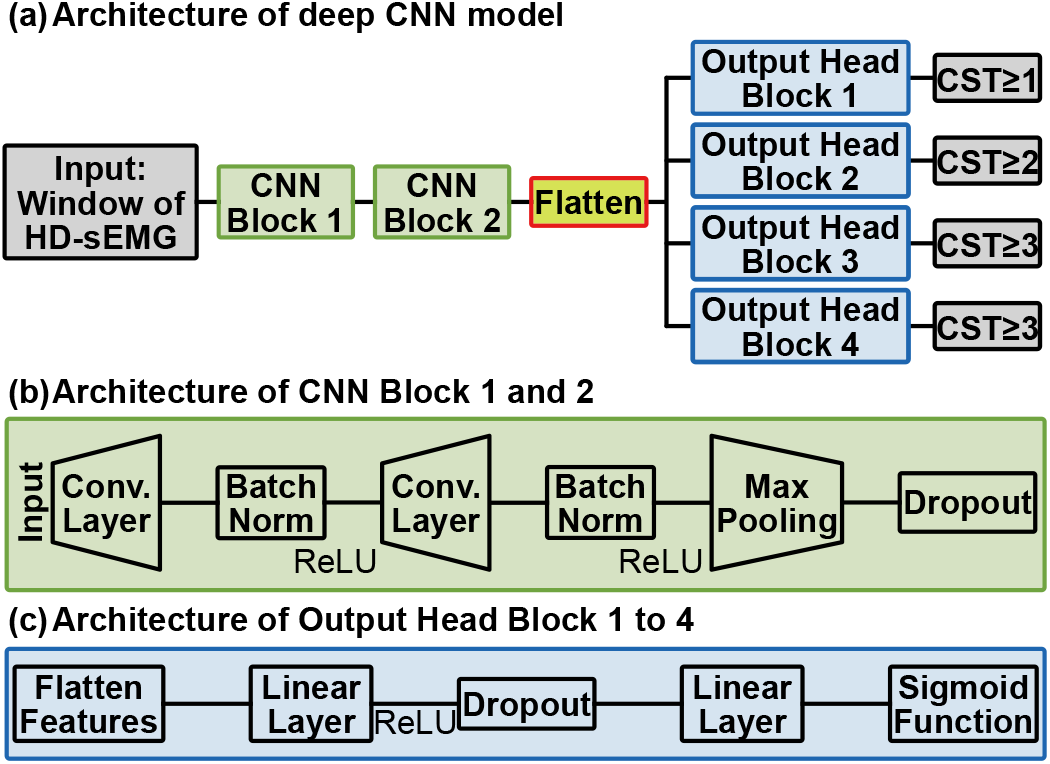
Architecture of the deep CNNs for neural drive decoding from HD-sEMG signals. All models consisted of two convolutional blocks, each comprising two convolutional layers, batch normalization, max pooling, and dropout. Features extracted from the convolutional blocks were flattened and passed to a multi-head dense architecture to decode neural drive as cumulative spike trains (CSTs). The 1D, 2D, and 3D CNNs shared the same network architecture and differed only in the dimensionality of the convolutional kernels, enabling the extraction of temporal, spatial, and spatiotemporal features, respectively.

To investigate the impact of convolutional kernel dimensionality on the generalizability of neural drive estimation, we built three versions of this model that share the same overall architecture but differ in dimensionality of their convolutional kernels (1D, 2D, or 3D); the same HD-sEMG signal was reshaped accordingly to match each kernel dimensionality.

The architectural hyperparameters of the three models are summarized in Table I. We kept the kernel size (3 along each dimension), and the activation functions identical across the three models; however, to offset the additional parameters and computational cost introduced by the higher-dimensional kernels, we reduced the number of kernel filters in each convolutional layer to one-half and one-quarter of those used in the 1D CNN for the 2D and 3D CNNs, respectively. As a result, the 1D, 2D, and 3D CNNs comprised 608708, 216228, and 563892 trainable parameters, respectively.

**TABLE I:**
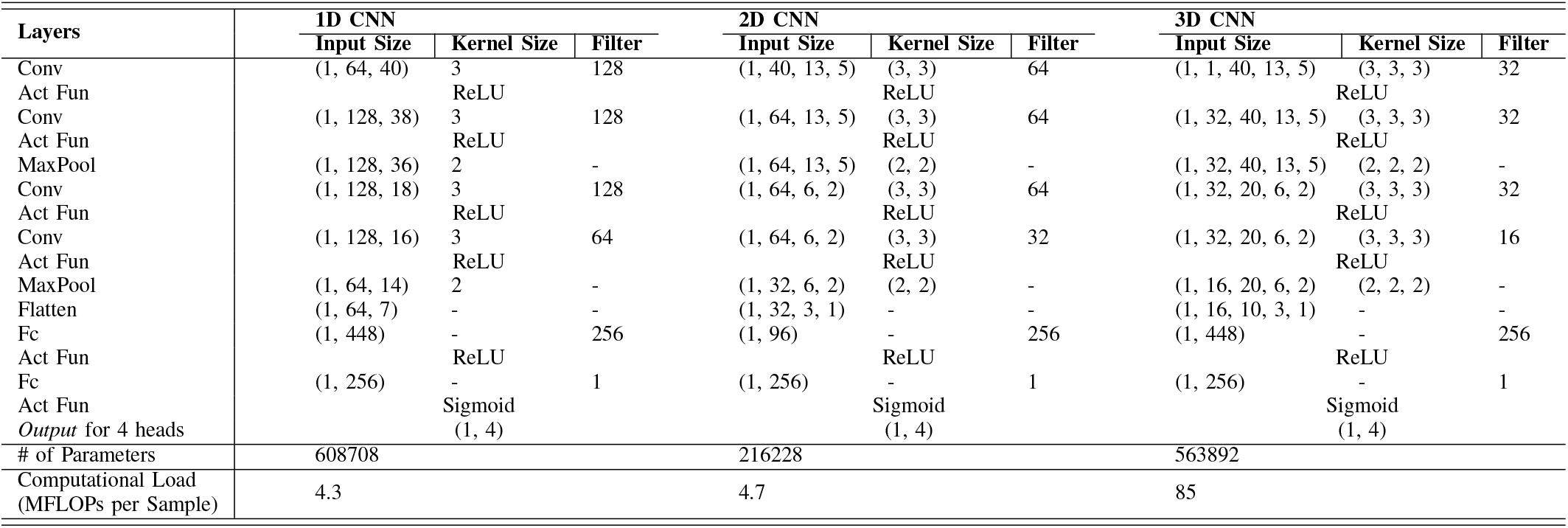
The structure of the 1D, 2D, and 3D CNNs used to decode neural drive from HD-sEMG signal. The abbreviations Conv, Act Fun, ReLU, MaxPool, and Fc denote a convolutional layer, Activation Function, Rectified Linear Unit, and a fully connected layer. Note that this table only includes one head of dense layers as all 4 heads are identical, and the output row indicates the concatenated dimension of the output from all 4 heads.

| Layers | 1D CNN |  |  | 2D CNN |  |  | 3D CNN |  |  |
| --- | --- | --- | --- | --- | --- | --- | --- | --- | --- |
|  | Input Size | Kernel Size | Filter | Input Size | Kernel Size | Filter | Input Size | Kernel Size | Filter |
| Conv | (1, 64, 40) | 3 | 128 | (1, 40, 13, 5) | (3, 3) | 64 | (1, 1, 40, 13, 5) | (3, 3, 3) | 32 |
| Act Fun |  | ReLU |  |  | ReLU |  |  | ReLU |  |
| Conv | (1, 128, 38) | 3 | 128 | (1, 64, 13, 5) | (3, 3) | 64 | (1, 32, 40, 13, 5) | (3, 3, 3) | 32 |
| Act Fun |  | ReLU |  |  | ReLU |  |  | ReLU |  |
| MaxPool | (1, 128, 36) | 2 | - | (1, 64, 13, 5) | (2, 2) | - | (1, 32, 40, 13, 5) | (2, 2, 2) | - |
| Conv | (1, 128, 18) | 3 | 128 | (1, 64, 6, 2) | (3, 3) | 64 | (1, 32, 20, 6, 2) | (3, 3, 3) | 32 |
| Act Fun |  | ReLU |  |  | ReLU |  |  | ReLU |  |
| Conv | (1, 128, 16) | 3 | 64 | (1, 64, 6, 2) | (3, 3) | 32 | (1, 32, 20, 6, 2) | (3, 3, 3) | 16 |
| Act Fun |  | ReLU |  |  | ReLU |  |  | ReLU |  |
| MaxPool | (1, 64, 14) | 2 | - | (1, 32, 6, 2) | (2, 2) | - | (1, 16, 20, 6, 2) | (2, 2, 2) | - |
| Flatten | (1, 64, 7) | - | - | (1, 32, 3, 1) | - | - | (1, 16, 10, 3, 1) | - | - |
| Fc | (1, 448) | - | 256 | (1, 96) | - | 256 | (1, 448) | - | 256 |
| Act Fun |  | ReLU |  |  | ReLU |  |  | ReLU |  |
| Fc | (1, 256) | - | 1 | (1, 256) | - | 1 | (1, 256) | - | 1 |
| Act Fun |  | Sigmoid |  |  | Sigmoid |  |  | Sigmoid |  |
| Output for 4 heads |  | (1, 4) |  |  | (1, 4) |  |  | (1, 4) |  |
| # of Parameters | 608708 |  |  | 216228 |  |  | 563892 |  |  |
| Computational Load (MFLOPs per Sample) | 4.3 |  |  | 4.7 |  |  | 85 |  |  |

### C. Data Processing

#### 1) CST “ground truth” from HD-sEMG decomposition

As shown in Fig.3, CST ground truth was obtained from the HDsEMG recordings through a standard decomposition pipeline consisting of band-pass filtering, bad-channel removal, and motor unit decomposition. Both the training and evaluation datasets underwent the same standard preprocessing prior to decomposition. The HD-sEMG signals were first band-pass filtered with a second-order Butterworth filter (cutoff frequencies of 20 and 500 Hz) to attenuate noise and motion artifacts [28]. The filtered signals were then visually inspected to identify channels with a low signal-to-noise ratio (SNR) or pronounced motion artifacts; these bad channels were replaced with the average of their neighboring channels to ensure the quality of the decomposed motor unit spike trains (MUSTs). Note that channel replacement was performed only for decomposition: the raw HD-sEMG signals were used for deep CNN training, validation, and testing.

**Fig. 3:**
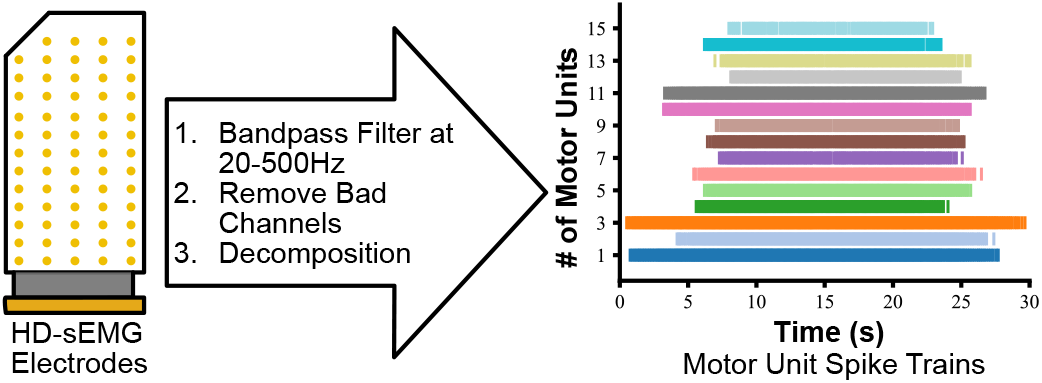
Pipeline to extract the ground-truth neural drive from HD-sEMG decomposition. The HD-sEMG signals were bandpass filtered to remove noise, and then decomposed by a blind source separation algorithm into the spike trains of individual motor units. The ground-truth neural drive was calculated as the CST, which is obtained by summing these motor unit spike trains.

For the training dataset, the preprocessed signals were decomposed into MUSTs using the blind source separation (BSS) algorithm of Negro et al. [8]. For the evaluation datasets, we instead used the pre-decomposed MUSTs provided with the data, which had been obtained using another BSS algorithm—convolutional kernel compensation [10]. Following decomposition, the resulting MUSTs were inspected, and only motor units with more than 100 spikes were retained [29]. A participant was excluded entirely if fewer than five valid motor units remained after applying this criterion in any of their recorded conditions (i.e., any session, muscle, or contraction intensity, as applicable to the given dataset). The CST ground truth was then obtained by summing the retained MUSTs at each time instant.

#### 2) Data preparation for deep CNN training and evaluation

The deep CNN models took the raw HD-sEMG signals as input and the corresponding decomposed CST as the training target. To construct the paired input–output samples, we followed the segmentation procedure of our previous work [16], [23]. We first aligned each spike index to the center of its MUAP, which was extracted from the decomposed motor unit spike trains. The aligned signals were then segmented with a sliding window (Fig. 4(a) and (b)). For the HD-sEMG input, each window spanned 40 data points with a step size of 20 data points, producing a 20-point (50%) overlap between consecutive windows. For the spike trains used to form the CST target, the window size and step size were both set to 20 data points; this non-overlapping segmentation tiles the signal so that each spike contributes to exactly one window and is never repeated across adjacent frames.

**Fig. 4:**
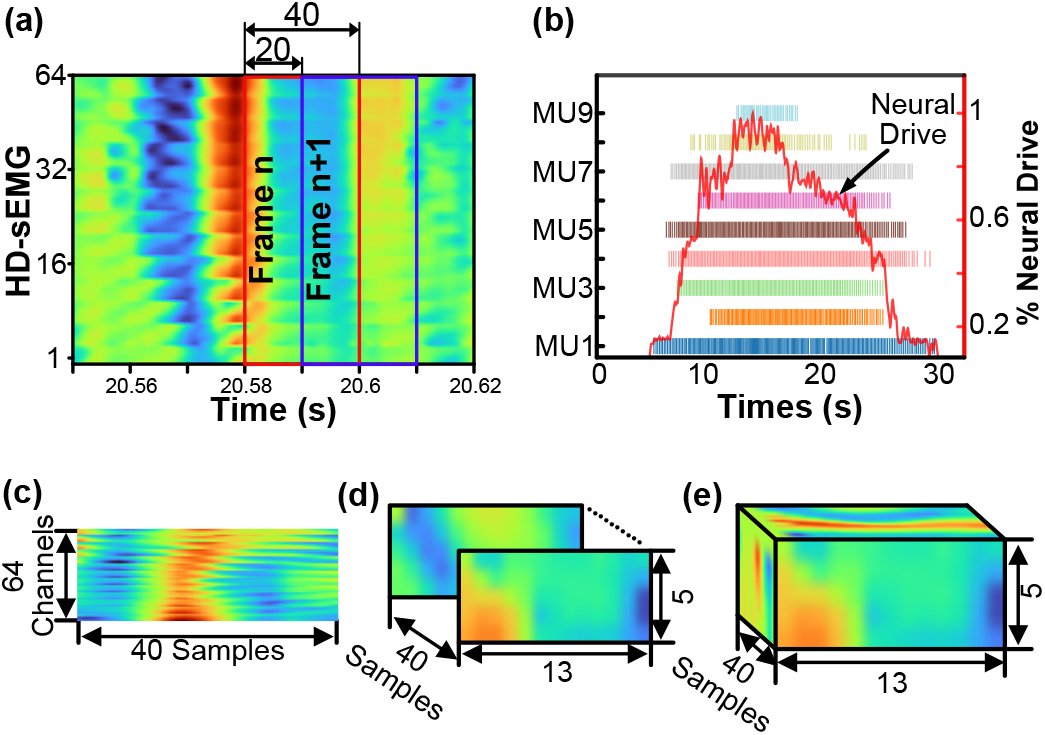
**(a)** Six consecutive frames of the flattened HD-sEMG signal in which the time duration is about 68.4 ms. The red window shows the *n*^*th*^ frame, which includes 40 samples, and the blue window shows the (*n* + 1)^*th*^ frame. The *n*^*th*^ and (*n* + 1)^*th*^ frame has 20 samples overlapped; **(b)** The output of the deep CNN, which was the neural drive represented as CST; **(c)** The shape of the HD-sEMG signal for 1D CNN; **(d)** The shape of the HD-sEMG after padding and reshaping for 2D CNN; **(e)** The shape of a HD-sEMG frame for 3D CNN.

We then reshaped each segmented 64×40 HD-sEMG frame into 13 × 5 × 40 to preserve the spatial features of HD-sEMG signals for both 2D and 3D CNNs (Fig. 4(d) and (e)). Because the HD-sEMG grid had one missing electrode at the top corner, a zero-valued channel was first added before the first channel of the original segmented HD-sEMG frame, expanding the input from 64 to 65 channels. The expanded 65 × 40 HDsEMG frame was then reshaped into 13 × 5 × 40, where the first two and the last dimension represented the spatial and temporal dimensions, respectively.

### D. Training and Validation of Deep CNNs

To train and validate the deep CNNs on the training dataset, we adopted a nested cross-validation scheme, as illustrated in Fig. 5. The training dataset comprised four recording conditions, formed by crossing the two muscles (VM and VL) with the two sessions recorded on separate days (day1 and day2)— namely day1 VM, day1 VL, day2 VM, and day2 VL—each containing three repeated contractions (C1–C3) from all five participants. The outer loop implemented four-fold leave-one-condition-out cross-validation, with each fold corresponding to one recording condition. Across the four runs, each condition was held out once as the test fold, while the remaining three conditions were used to train and validate the model. Within each run, the inner loop applied a leave-one-contraction-out scheme to further split three training conditions: two of the three contractions were used to train the CNN, and the remaining contraction served as the validation set during training. This inner split was rotated three times so that each contraction was used once for validation, yielding three subfolds and thus three trained models per run. Each of these three models was then evaluated on the unseen held-out condition. Repeating this procedure across the four outer folds produced a total of 4 × 3 = 12 trained models.

**Fig. 5:**
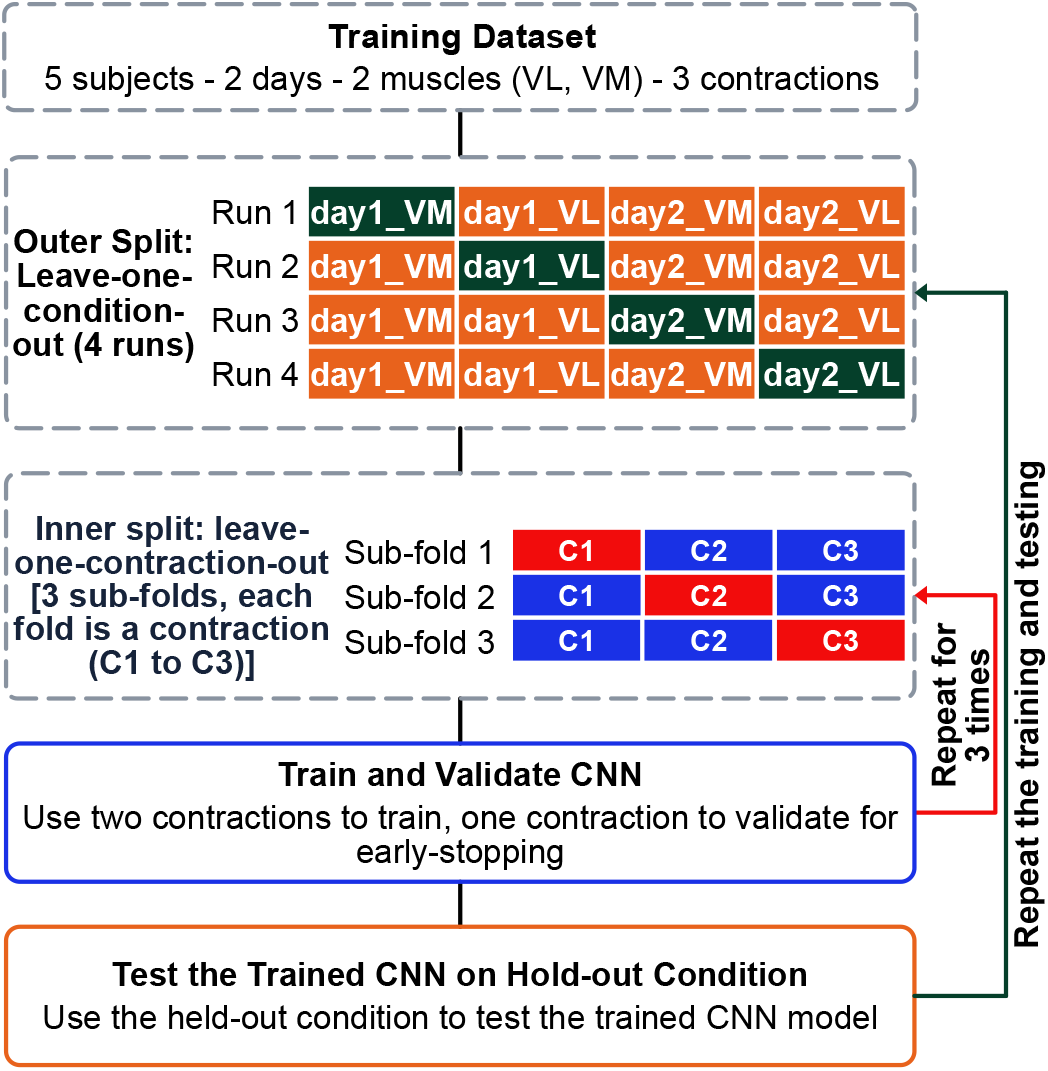
Nested cross-validation pipeline for training and validating the deep CNNs on the training dataset (5 subjects, 2 days, 2 muscles, 3 contractions). **Outer split:** a leave-one-condition-out scheme over the four day × muscle conditions, across four runs, each condition is held out once as the test set (green) while the remaining three form the training set (orange). **Inner split:** within each run, the three contractions (C1–C3) of the training conditions are divided by leave-one-contraction-out into three sub-folds, each sub-fold using two contractions for training (blue) and one for validating the model (red).

All CNN models were implemented, trained, and evaluated in Python using the PyTorch deep learning framework. Training was performed using a learning rate of 0.001 and optimized with the Root Mean Square Propagation (RMSProp) optimizer. Binary cross-entropy was used as the loss function. For all deep CNNs, training was performed for up to 200 epochs. An early-stopping scheme was applied to terminate training when the change in the validation F1 score remained below 0.001 for 50 consecutive epochs. The CNNs were trained on a workstation equipped with dedicated GPUs. After training, model evaluations were conducted on a laptop equipped with an NVIDIA RTX 3000 GPU and an Intel Ultra 7 155H CPU to assess the generalizability and computational efficiency of the trained deep CNN models.

### E. Evaluating the Generalizability of the Trained CNNs

#### 1) Evaluating the trained CNNs with the cross-intensity evaluation dataset

After training the CNNs, we first evaluated the models on the entire cross-intensity evaluation dataset, estimating neural drive for each subset. Because this dataset was recorded across different intensities, its primary purpose was to assess the cross-intensity generalizability of the trained CNNs. In addition, it was recorded from participants who were not part of the training dataset, allowing us to evaluate cross-participant generalizability.

#### 2) Evaluating the trained CNNs with the cross-muscle evaluation dataset

We also evaluated the trained models on the entire cross-muscle evaluation dataset, estimating neural drive for each subset. Because this dataset was recorded across different muscles, its primary objective was to assess the cross-muscle generalizability of the trained CNNs. In addition, like the cross-intensity dataset, it was recorded from different participants of the training dataset, allowing us to also evaluate cross-participant generalizability.

### F. Evaluation Metrics

As the effective neural drive that produces muscle force lies in the low-frequency band [30], [31], we smoothed the estimated CSTs with a 400-ms Hanning-window filter to obtain neural drive [6]. Then we performed a regression analysis, following the same procedure as in the NMI application [15], [32]. Specifically, we tuned a linear regression equation without a bias term, using the neural drive estimated by the CNNs as the input (**x**) and the neural drive estimated by the BSS method as the target (**y**), and solved for the slope *α* by the least squares method,

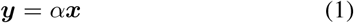

where ***x*** is the smoothed CST estimated by the CNNs and ***y*** is the smoothed CST estimated by the BSS method. We then used the fitted *α* to compute the mapped neural drive (***ŷ***) from the CNN-estimated neural drive, and calculated the correlation coefficient (*R*) and the normalized root mean square error (nRMSE) between the mapped neural drive and the BSS-estimated neural drive, as defined below.

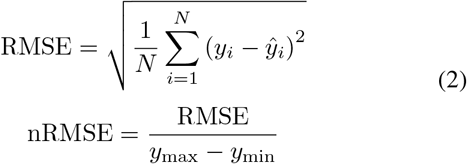

where *y*_*i*_ is the neural drive estimated by the BSS method, *ŷ*_*i*_ is the neural drive mapped from the CNN-estimated neural drive, *N* is the total number of samples in the estimated neural drive, and *y*_max_ and *y*_min_ are the maximum and minimum values of the BSS-estimated neural drive.

Finally, we evaluated the computational efficiency of the three CNN models by measuring their average per-sample inference time on both a GPU and a CPU. We randomly selected 500 segmented HD-sEMG signals from the evaluation datasets and used each model to estimate the CST for each sample, recording the elapsed prediction time. For each CNN model—computational device combination, we repeated this procedure for 20 times to exclude any random factor of the CPU or GPU devices, and used the averaged inference time per sample as the measured inference time.

### G. Statistical Analysis

All statistical analyses were performed in R (version 4.5.3). To assess the impact of convolutional kernel dimensionality on the generalizability of the CNN models, we fitted a linear mixed-effects model (*lme4::lmer*) of the form

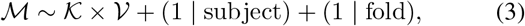

where the response ℳ is a generalizability metric, modeled separately for *R* and nRMSE (i.e., a separate model was fitted for each metric). The fixed effects are the dimensionality of the convolutional kernel (*K*) and the evaluation scenario, which includes cross-intensity and cross-muscle (*V*); the term *K* × *V* denotes both main effects and their interaction. The model included two crossed random intercepts: (1 | subject), which accounts for subject-specific variability in the evaluation dataset [33], and (1 | fold), which accounts for variability across the leave-one-condition-out cross-validation folds.

The significance of each fixed effect and of their interaction was evaluated using a two-way ANOVA test on the fitted model (*lmer Test*). We used *p* < 0.05 as the criterion for statistical significance. If statistical significance was determined for the interaction between fixed effects, we would compute Tukey-adjusted pairwise comparisons of estimated marginal means (*emmeans::emmeans*) for the interaction effects, i.e., differences between the levels of one factor within each level of the other.

## III Results

### A. Generalizability of Deep CNNs Evaluated by Cross-Intensity Dataset

As shown in Fig. 6(a), all three CNN models accurately estimated the neural drive and showed strong agreement with the ground-truth neural drive on the cross-intensity dataset, which was recorded across three contraction intensities from a participant group independent of the training dataset. The two-way ANOVA on the fitted LMM did not detect a significant interaction effect between kernel dimensionality and contraction intensity for *R*. Among the three models, the 2D CNN achieved the highest overall *R* (0.988 ± 0.006) across all intensities, while the 3D and 1D CNNs showed their lowest mean *R* at 10% and 50% MVC, respectively. These two models were also the most sensitive to inter-participant variation, but at opposite intensities: the 3D CNN was most variable at 50% MVC and the 1D CNN at 10% MVC (standard deviation 0.02 and 0.01, respectively).

**Fig. 6:**
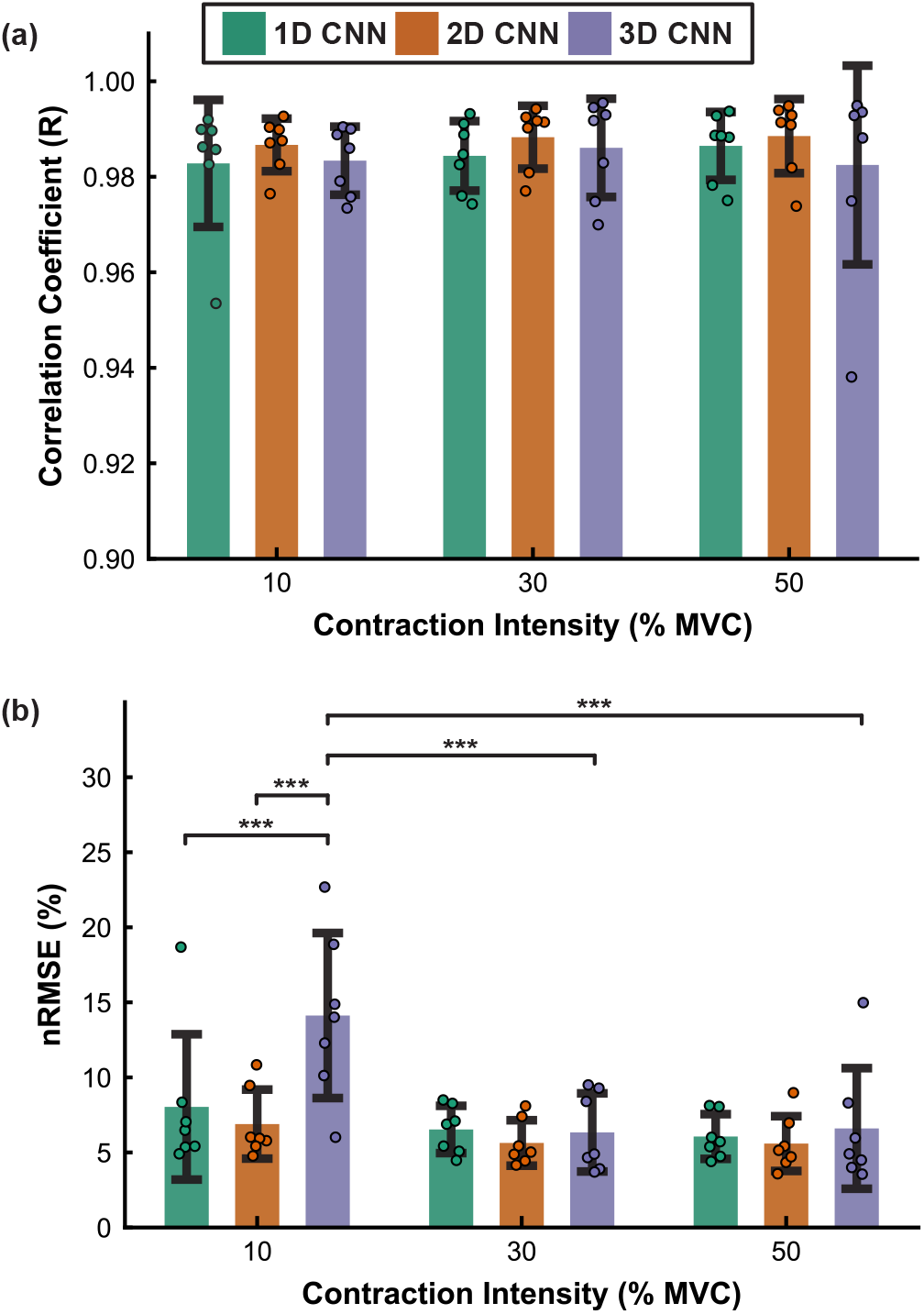
Generalizability of the deep CNNs evaluated on the cross-intensity dataset (recorded at 10%, 30%, and 50% MVC). (a) Correlation coefficient (*R*) and (b) normalized root-mean-square error (nRMSE) between the BSS-estimated neural drive and the neural drive mapped from the CNN-estimated neural drive.

In contrast, nRMSE depended more strongly on both factors (Fig. 6(b)). The two-way ANOVA detected significant main effects of kernel dimensionality (*p* < 0.001) and contraction intensity (*p* < 0.001), as well as a significant interaction between them (*p* < 0.001). Post hoc comparisons localized this interaction to the lowest intensity: at 10% MVC, the 3D CNN estimated the neural drive with significantly higher nRMSE than both the 1D and 2D CNNs (*p* < 0.001), whereas the three models did not differ significantly at 30% or 50% MVC. Examined within each model, nRMSE at 10% MVC was significantly higher than at the other intensities for the 3D CNN (*p* < 0.001), whereas the 1D and 2D CNNs were more robust to inter-intensity variation. These trends are consistent with the averaged linear-fit slope *α* in Table II: the 3D CNN had the lowest *α* at 10% MVC, indicating that, under this condition, the amplitude of the neural drive is estimated to have deviated most from the BSS-estimated neural drive.

**TABLE II:** Averaged linear-fit slope *α* between the CNN- and BSS-derived neural drive for the 1D, 2D, and 3D CNNs at each contraction intensity of the cross-intensity dataset.

|  | 10%MVC | 30%MVC | 50%MVC |
| --- | --- | --- | --- |
| 1D CNN | $0.71 \pm 0.17$ | $0.86 \pm 0.24$ | $0.77 \pm 0.18$ |
| 2D CNN | $0.77 \pm 0.17$ | $0.88 \pm 0.21$ | $0.81 \pm 0.20$ |
| 3D CNN | $0.65 \pm 0.20$ | $0.79 \pm 0.22$ | $0.73 \pm 0.18$ |

### B. Generalizability of Deep CNNs Evaluated by Cross-Muscle Dataset

As shown in Fig. 7(a), on the cross-muscle dataset—recorded across three muscles from a participant group independent of the training data—*R* was governed primarily by the recorded muscle rather than by the convolutional kernel dimensionality. The two-way ANOVA on the fitted LMM detected a significant main effect of muscle (*p* < 0.001): the GL muscle yielded the lowest and most variable *R*, whereas the GM and SOL muscles yielded higher and mutually comparable values. The 1D CNN achieved the lowest *R* across muscles, whereas the 2D and 3D CNNs achieved comparable but higher values.

**Fig. 7:**
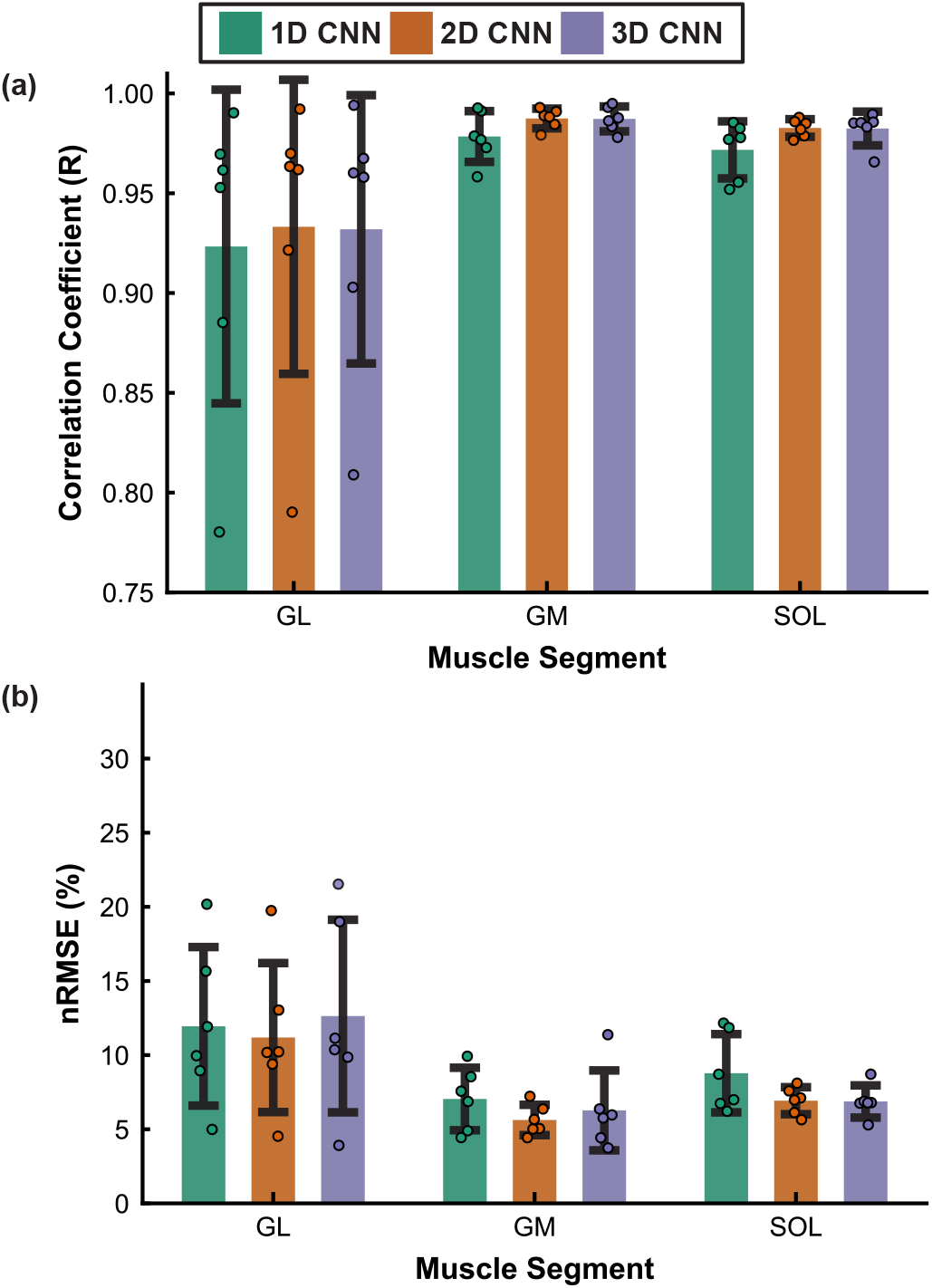
Cross-muscle generalizability of the 1D, 2D, and 3D CNNs evaluated on the independent testing dataset (recorded at GM, GL and SOL muscles). (a) Correlation coefficient and (b) normalized root-mean-square error between the BSS-estimated neural drive and the neural drive mapped from the CNN-estimated neural drive, shown for each muscle.

A similar pattern was observed for nRMSE (Fig. 7(b)). The two-way ANOVA revealed a significant main effect of muscle (*p* < 0.001). Among the three muscles, GL exhibited the highest and most variable nRMSE, GM the lowest, and SOL intermediate values. For both GM and SOL, the 2D and 3D CNNs achieved lower nRMSE than the 1D CNN. The averaged linear-fit slope *α* (Table III) generally reflected the trends observed in nRMSE for GM and GL, with models producing larger nRMSE also exhibiting smaller *α* values. However, a different pattern was observed for SOL: although its average nRMSE was comparable to GM’s, its *α* was approximately 40% lower. This discrepancy suggests that the CNN models produced a substantial number of false-positive (FP) predictions in SOL. Because nRMSE quantifies the overall magnitude difference between the estimated and reference neural drives, these FP predictions had a limited impact on nRMSE, but they reduced the slope of the linear relationship, thereby lowering *α*.

**TABLE III:** Averaged linear-fit slope *α* between the CNN- and BSS-derived neural drive for the 1D, 2D, and 3D CNNs at each muscle of the cross-muscle dataset.

|  | GL | GM | SOL |
| --- | --- | --- | --- |
| 1DCNN | $0.33 \pm 0.21$ | $0.60 \pm 0.31$ | $0.23 \pm 0.11$ |
| 2DCNN | $0.43 \pm 0.22$ | $0.67 \pm 0.32$ | $0.31 \pm 0.17$ |
| 3DCNN | $0.41 \pm 0.22$ | $0.64 \pm 0.31$ | $0.35 \pm 0.21$ |

Further inspection showed that the reductions in *R* and increases in nRMSE observed for the GL muscle were driven by only two of the six subjects, regardless of the CNN architecture. To investigate this finding, we plotted the neural drive estimated by BSS and the 2D CNN models for one of these subjects. As shown in Fig. 8, the neural drive estimated by BSS was largely decoupled from the measured contraction-intensity trajectory of the trapezoidal shape. Instead of plateauing at the sustained-intensity output, the neural drive remained relatively low for approximately the first 70% of the contraction plateau, then increased sharply near the end of the contraction. However, the neural drive estimated by all CNN models continued to follow the overall intensity profile.

**Fig. 8:**
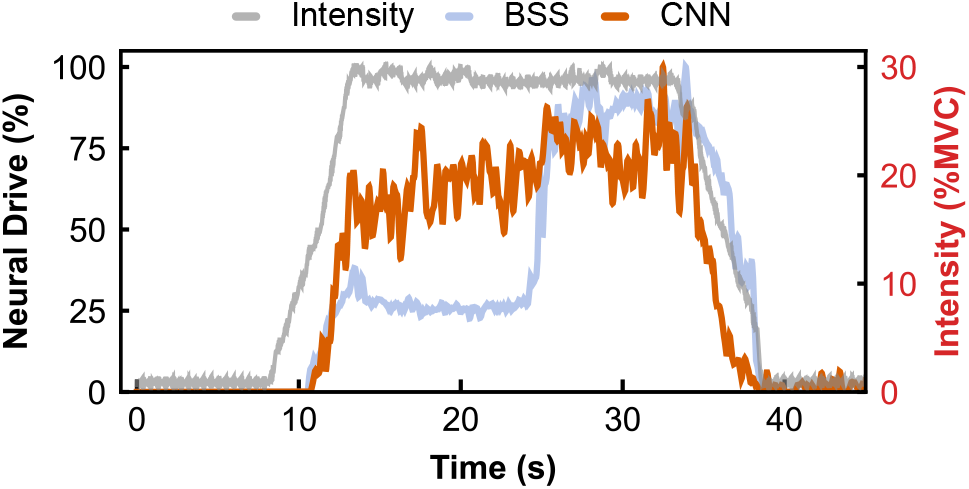
The representative neural drive estimated by the BSS method from HD-sEMG data collected from the GL muscle for one of the outlier subjects corresponds to the measured contraction intensity.

### C. Computational Efficiency of Different CNN Architectures

As shown in Fig. 9, inference time depended on both the kernel dimensionality and the computing platform. On the CPU, inference time increased significantly with kernel dimensionality: the 3D CNN was the slowest at 2.11 ± 0.05 ms/sample, roughly 3.5× slower than the 1D (0.62 ± 0.06 ms/sample) and 2D (0.66 ± 0.05 ms/sample) CNNs, which were comparable. Moving from the CPU to the GPU reduced the inference time of the 3D CNN by approximately 32% (to 1.42 ± 0.08 ms/sample), but increased that of the 1D and 2D CNNs, with the 1D CNN rising the most—by roughly 130%, to 1.43 ± 0.08 ms/sample. As a result, on the GPU, the inference times of the three CNN architectures converged to similar values, with 1D and 2D CNNs no longer much faster than 3D CNNs. Additionally, the inference time of the 2D CNN on the GPU is marginally faster than that of the 1D CNN (10% faster, 1.27 ± 0.04)

**Fig. 9:**
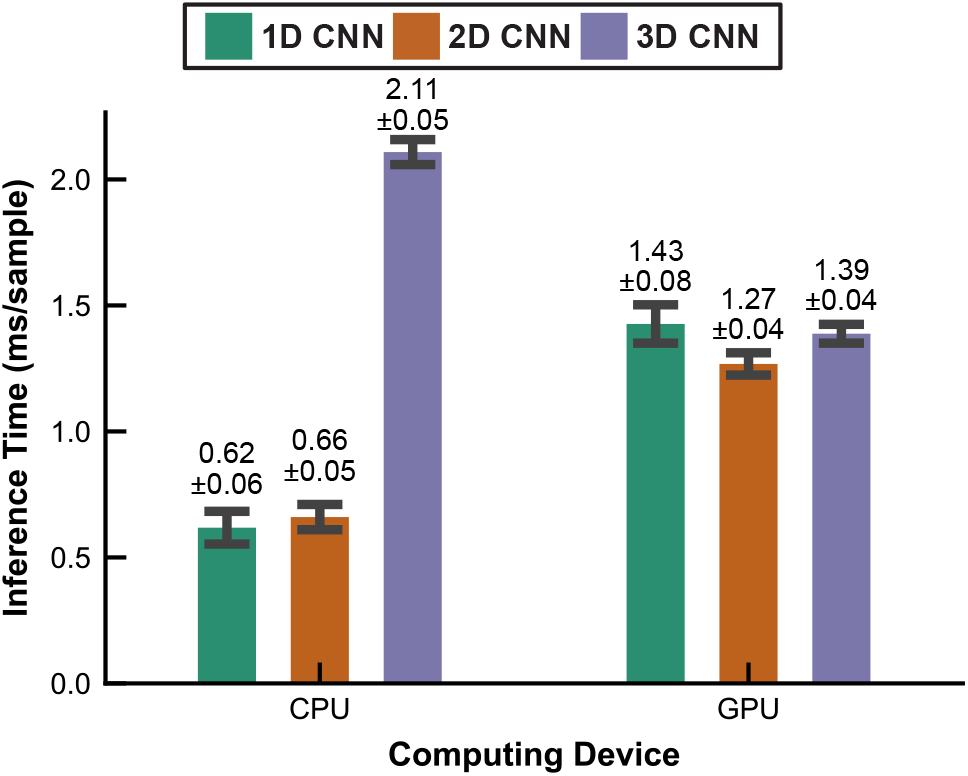
Average inference time per sample for the 1D, 2D, and 3D CNNs on CPU and CUDA-enabled GPU platforms. Bars and error bars denote the mean and standard deviation. On the CPU, inference time increased sharply with kernel dimensionality, the 3D CNN being the slowest; on the GPU, the three models converged to a similar range, as GPU execution reduced the cost of the 3D CNN but increased that of the 1D and 2D CNNs.

## IV. Discussion

This study investigated how the dimensionality of CNN kernels affects the generalizability and computational efficiency of neural drive estimation from HD-sEMG. The CNNs were trained on a dataset that included multiple participants and muscles at a single intensity across two days. We evaluated the generalizability of trained CNNs on two unseen datasets collected from separate groups of participants—one across three contraction intensities and the other across three muscles. Without any retraining, all three models generalized to both unseen datasets, but kernel dimensionality affected performance only under limited conditions (Fig. 6 and Fig. 7). In contrast, computational efficiency depended strongly on kernel dimensionality and computing platform (Fig. 9): the 3D CNN was markedly slower than the 1D and 2D CNNs on the CPU, whereas all three models achieved comparable inference times on the GPU.

Contrary to our hypothesis, increasing kernel dimensionality did not improve generalizability. Although the 3D CNN was expected to perform best by exploiting spatiotemporal MUAP features, it performed comparably to the 2D CNN, and both models outperformed the 1D CNN only marginally across the evaluation datasets (Fig. 6 and Fig. 7). A potential explanation lies in the feature-extraction mechanism of 3D convolution: as originally proposed, 3D kernels capture the temporal evolution of spatial features across consecutive video frames [34]. This corresponds to modeling how spatial MUAP features evolve over time across the HD-sEMG electrode grid. However, because all datasets were collected during isometric contractions, in which muscle length remained relatively unchanged, the spatial MUAP features across the HD-sEMG grid likely changed little over time. As a result, explicitly modeling the temporal evolution of these spatial features may have provided limited additional benefit beyond that offered by 2D spatial kernels. This interpretation is consistent with prior HD-sEMG gesture-recognition work showing that 3D CNNs provide only modest gains for static gestures but larger benefits for dynamic movements [35], [36]. Whether similar advantages can be realized for neural drive estimation during dynamic movements remains an open question and requires further investigation [23], [37].

When evaluated on the cross-intensity dataset, all three CNNs yielded the highest nRMSE at 10% MVC relative to 30% and 50% MVC (Fig. 6(b)). We attribute this to the greater relative contribution of noise in HD-sEMG signals recorded at low intensity, where fewer active motor units and lower firing rates reduce MUAP amplitude, thereby increasing the contribution of noise [38]. Because we used raw HD-sEMG signals as inputs, this noise-related contamination makes neural-drive estimation more challenging across all three architectures, consistent with previous observations in [39]. Moreover, the 3D CNN was the most sensitive model at low intensity. Its nRMSE was significantly higher than that of the 1D and 2D CNNs on the 10% MVC subset, and was also significantly higher than its own nRMSE at 30% and 50% MVC. Notably, as shown in Table I, the 3D CNN contained fewer trainable parameters than the 1D CNN (563,892 versus 608,708), suggesting that the observed degradation is unlikely to be explained solely by model size or over-parameterization [40]. Instead, the greater sensitivity of the 3D CNN was likely due to the degradation of spatiotemporal MUAP features at low intensities. Because 3D convolutional kernels capture the temporal evolution of spatial MUAP features, distortions in these weaker MUAP representations may disproportionately affect the 3D CNN relative to 1D and 2D CNNs. Nevertheless, the precise mechanism underlying the greater resilience of the 1D and 2D CNNs under low-intensity conditions remains unclear and requires further investigation. Future work could also examine whether transfer learning or fine-tuning with a small amount of target-domain data can mitigate this limitation, as suggested by a recent study [21].

Beyond kernel dimensionality, generalizability was also limited by atypical neural-drive patterns in the evaluation dataset, as shown in Fig 7 and 8. This decoupling of neural drive estimated by BSS and measured contraction intensity pattern may be attributed to several factors, including motor unit rotation [41] and muscle fatigue [42]. During motor unit rotation, some newly recruited or rotated motor units may not be fully identified by BSS methods, resulting in the omission of motor unit spike trains and a less complete representation of the underlying neural drive. Notably, the nRMSE values observed in the present study were comparable to those reported in a previous HD-sEMG-based force estimation study conducted under muscle fatigue conditions, where fatigue was shown to significantly impair estimation accuracy [43]. This observation suggests that muscle fatigue may be another plausible contributor to the decoupling observed in the affected subjects. Together, these findings highlight the need to further investigate the mechanisms underlying the misalignment between the neural drive derived by BSS and the trajectory of contraction intensity. In particular, future studies are needed to determine the respective contributions of muscle fatigue and motor unit rotation to this misalignment, as well as their impact on the generalizability of deep CNNs.

We also observed that the averaged linear-fit slope *α* (Tables II and III) was consistently below 1 across all models, muscles, and intensities, indicating that the CNNs estimated a larger number of spikes—and therefore a higher cumulative spike train amplitude—than the BSS reference. Our previous work [44] also found that the BSS algorithm tends to identify fewer motor unit spike trains in HD-sEMG data. These findings highlight an opportunity to further investigate the factors contributing to the observed misalignment between the CNN-estimated neural drive and the BSS-derived reference. Specifically, future studies are needed to determine whether the misalignment arises from false-positive detections by the CNN or motor unit discharges not identified by the BSS algorithm. Despite these findings, several limitations should be considered when interpreting the results. First, because the deep CNN models were trained and evaluated using neural drive derived from BSS algorithm rather than the true physiological neural drive, the findings should be interpreted as evidence of the models’ applicability to real-world NMI systems, rather than as insights into the physiological properties of neural drive. Second, the HD-sEMG dataset used for both training and evaluation was collected solely during lower-limb isometric contractions. Consequently, the proposed model’s ability to generalize beyond the lower-limb muscles remains unknown. Third, the evaluation dataset was limited in both the number of participants and population diversity. Therefore, although the proposed CNN models demonstrated promising generalizability within the studied cohort, their applicability to broader populations remains to be validated. Finally, our inference-time benchmarks were obtained on a laptop-class CPU and GPU rather than on the embedded processors typical of wearable NMI hardware, so the computational-efficiency findings should be regarded as indicative and verified on target hardware before deployment.

## V. Conclusion

This study systematically investigated the impact of convolutional kernel dimensionality on the generalizability and computational efficiency of deep CNNs for neural drive estimation from HD-sEMG signals. All three CNN architectures are generalized to two independent unseen HD-sEMG datasets collected under different participants, sessions, contraction intensities, and muscles, demonstrating the potential of CNN-based neural drive estimation to transfer across datasets without retraining. Although the 2D and 3D CNNs marginally outperformed the 1D CNN, the 3D CNN offered no meaningful advantage over the 2D CNN despite its higher CPU computational cost. These findings suggest that increasing architectural complexity beyond spatial convolution does not improve generalizability for neural drive estimation. Overall, the 2D CNN offered the best balance between estimation accuracy and computational efficiency, making it a practical choice for a deployable CNN-based neural drive estimator, particularly on resource-constrained platforms. Future work should expand the evaluation dataset to include larger, more diverse populations, additional muscle groups, and dynamic contractions, and further investigate the factors that limit the generalizability of deep CNN models for neural drive estimation.

